# Quantifying RNA Synthesis at Rate-Limiting Steps of Transcription Using Nascent RNA-Sequencing Data

**DOI:** 10.1101/2021.08.03.454856

**Authors:** Adelina Rabenius, Sajitha Chandrakumaran, Lea Sistonen, Anniina Vihervaara

## Abstract

Nascent RNA-sequencing tracks transcription at nucleotide resolution. The genomic distribution of engaged transcription complexes, in turn, uncovers functional genomic regions. Here, we provide data-analytical steps to 1) identify transcribed regulatory elements *de novo* genome-wide, 2) quantify engaged transcription complexes at enhancers, promoter-proximal regions, divergent transcripts, gene bodies and termination windows, and 3) measure distribution of transcription machineries and regulatory proteins across functional genomic regions. This protocol follows RNA synthesis and genome-regulation in mammals, as demonstrated in human K562 erythroleukemia cells.

For complete details on the use and execution of this protocol, please refer to Vihervaara, *et al.*, 2021.

**Graphical Abstract:** 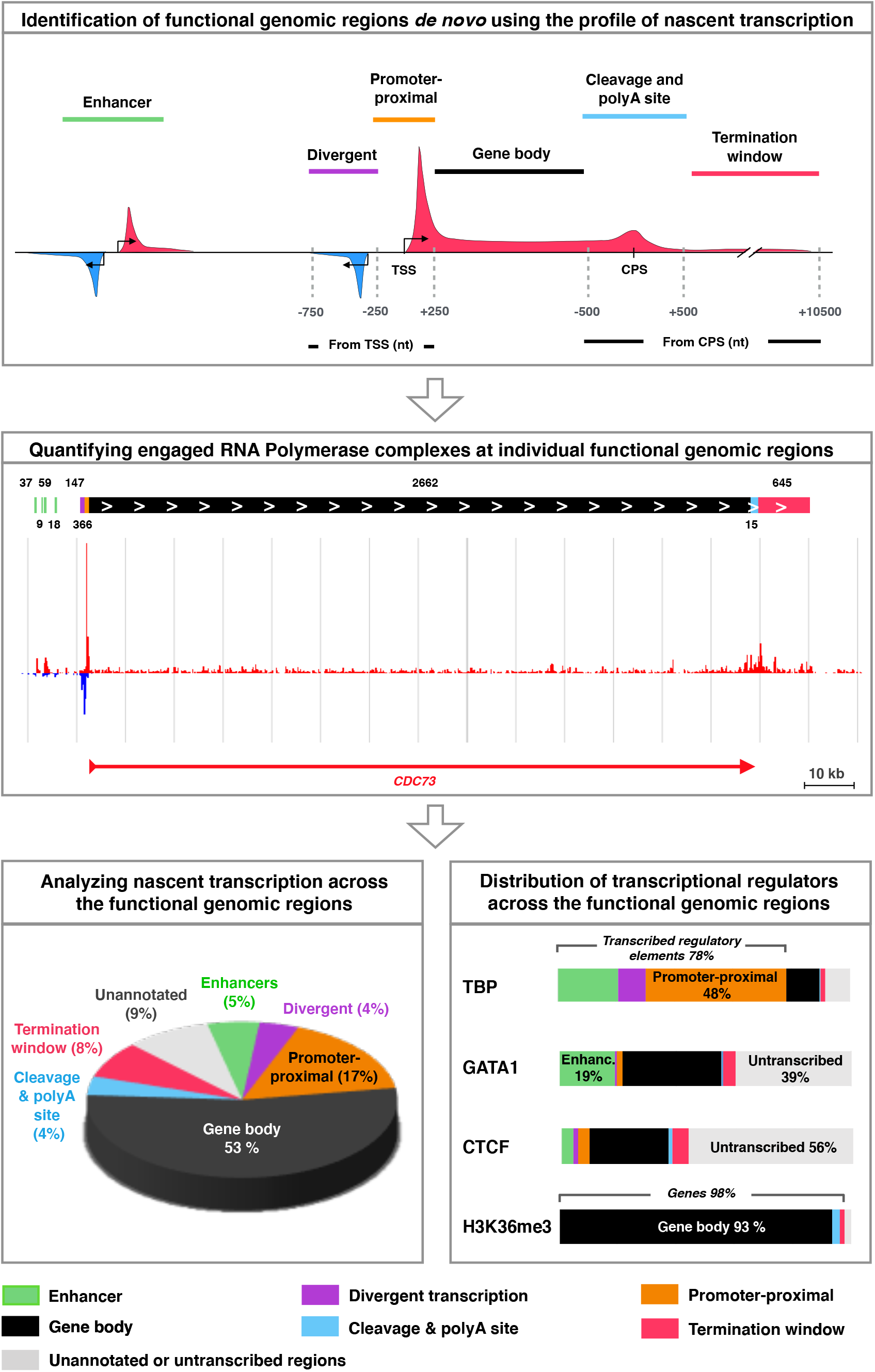

## Before You Begin

### Nascent transcription reveals functional genomic regions

Transcription is a fundamental process in every organism, and its coordination defines RNA synthesis in the cell. The process of RNA synthesis is orchestrated by a plethora of transcription factors, cofactors, chromatin remodelers and RNA-processing complexes (reviewed in Wissink *et al.*, 2019). The transcriptional regulators interact, directly or indirectly, with functional genomic regions, including promoters, enhancers, gene bodies and termination windows. Each cell executes a transcription program that reflects the cellular identity, cellular functions and the intra- and extracellular signaling pathways (reviewed in Zeitlinger and Stark, 2010; Vihervaara *et al.*, 2018). Consequently, the repertoire of active genes and regulatory elements differs in distinct cells, and the activity of functional genomic regions can rapidly adapt to changes in intra- and extra-cellular conditions.

Enhancers are distal regulatory elements that generally produce short and unstable transcripts called enhancer RNAs (eRNAs; Kim *et al.*, 2010). Enhancer transcription has been shown to precede activation of genes (Kim *et al.*, 2010; Kaikkonen *et al.*, 2013; Arner *et al.*, 2015), and eRNA production moderately correlates with the target genes’ activity (Henriques *et al.*, 2018; Mikhaylichenko *et al.*, 2018). Intriguingly, promoters and enhancers display a unified transcriptional architecture where two core initiation regions drive transcription into opposing directions (Core *et al.*, 2008; 2014, reviewed in Andersson and Sandelin, 2020). This pattern of divergent transcription conveys function to enhancers (Tippens *et al.*, 2020), and it allows identification of transcribed regulatory elements *de novo* (Danko et al, 2015; Azofeifa and Dowell, 2017; Vihervaara *et al.*, 2017; Chu *et al.*, 2018, Wang *et al.*, 2019).

Transcriptional activity from genes and enhancers is controlled at the level of local chromatin environment, recruitment and progression of RNA Polymerase II (Pol II) complexes, and the three-dimensional architecture of the linear DNA molecules (reviewed in Field and Adelman, 2020; Schoenfelder and Fraser, 2020). The regulatory signals at the chromatin culminate at rate-limiting steps of transcription (Figure 1), constituting of 1) chromatin opening, 2) assembly of the Pre-Initiation Complex, 3) initiation of transcription, 4) promoter-proximal Pol II pausing, 5) promoter-proximal Pol II pause-release, 6) productive elongation, 7) co-transcriptional RNA processing, 8) transcript cleavage, 9) transcription termination, and 10) recycling of Pol II (reviewed in Fuda *et al.*, 2009; Wissink *et al.*, 2019). The rate-limiting steps of transcription occur at distinct genomic regions (Figure 1). Consequently, assays that track nascent transcription in a population of cells, display characteristic ‘signatures’ of engaged Pol II molecules at transcribed enhancers, active promoters, transcription start sites (TSSs), promoter-proximal pause-regions, gene bodies, cleavage and polyadenylation sites (CPSs), and termination windows (Figure 1B-D). Here, we provide data-analytical steps to identify transcriptionally active functional genomic regions in the investigated condition. The nucleotide-resolution power of nascent RNA sequencing is further harnessed to count engaged Pol II complexes at each identified functional genomic region. Quantifying engaged Pol II complexes and transcriptional regulators at distinct genomic regions provides a framework for detailed analyses of nascent transcription and its regulation.

**Figure 1.**
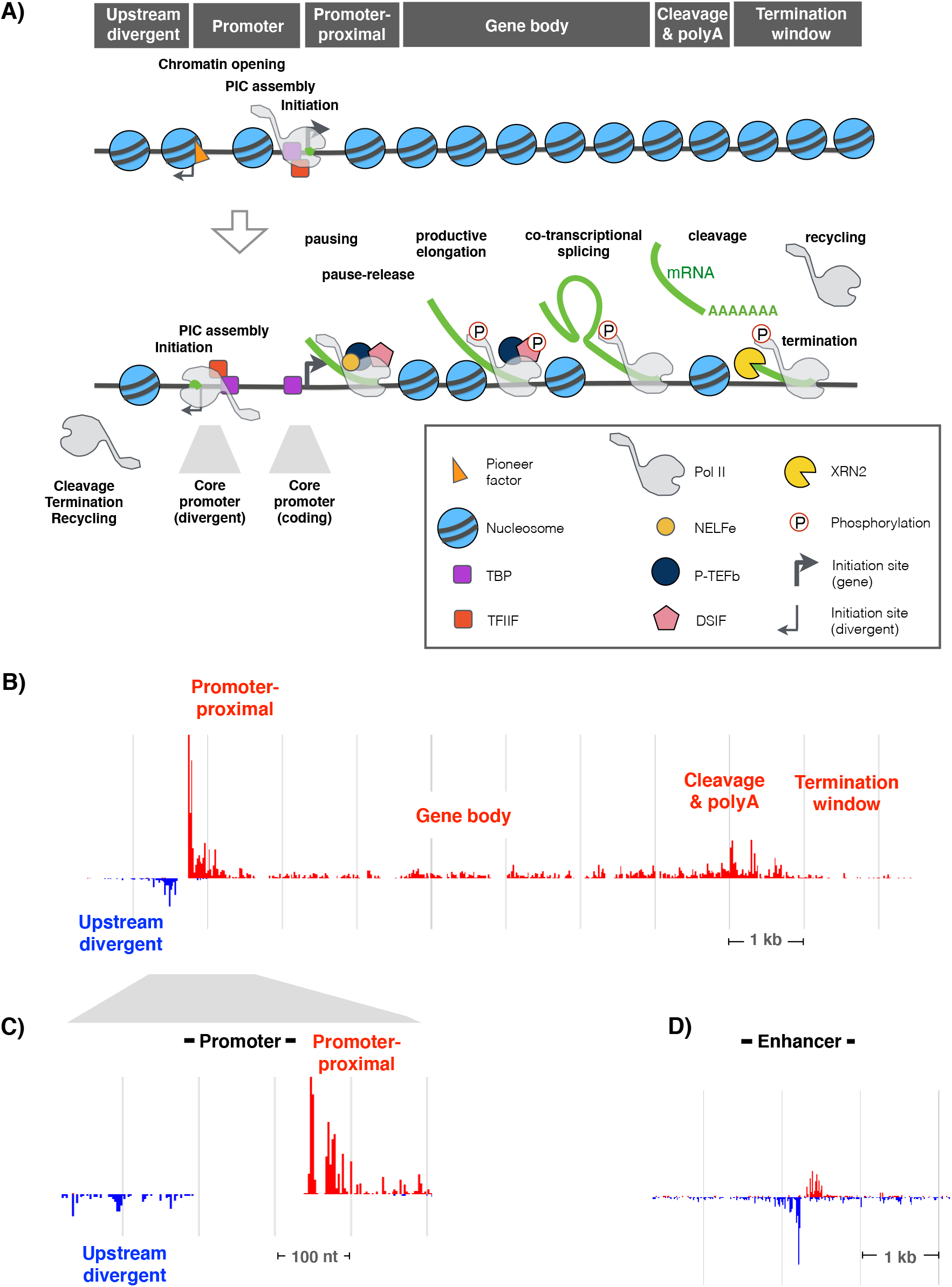
Steps of transcription occur at distinct functional genomic regions. **A)** Schematic presentation of rate-limiting steps of transcription. 1) Closed chromatin is opened by pioneer transcription factors (orange triangle) and chromatin remodellers. 2) Pre-initiation complex (PIC), including TBP and TFIIF, assembles at the core promoters and positions Pol II to initiation. 3) Pol II initiates transcription, 4) pauses at the promoter-proximal region, and 5) is released by P-TEFb into 6) productive elongation. 7) Co-transcriptional processes, such as 5’-capping and splicing, occur while Pol II progresses through the gene. At the end of the gene, 8) the transcript is cleaved and polyadenylated (polyA). 9) XRN2 degrades the uncapped transcript, chasing Pol II and causing transcription to terminate. 10) Pol II is released from the chromatin and recycled into new initiation. **B)** Profile of nascent transcription at the distinct regions of a typical gene in humans. Transcription profile at **C)** promoter and **D)** enhancer. The gene in B and C is *RPSA*, and the enhancer in D is one of the functionally verified enhancers of *beta*-globin locus control element.

### Obtaining datasets of nascent RNA sequencing

This protocol relies on strand-specific mapping of RNA synthesis. The strand-specificity is essential for identification of active promoters and enhancers *de novo* based on their pattern of divergent transcription (Core *et al.*, 2008; 2014; Danko et al, 2015; Seila *et al.*, 2008; Tippens *et al.*, 2018; 2020; Vihervaara *et al.*, 2017; Wang *et al.*, 2019). Besides strand-specific mapping of RNA synthesis, the precise positions of active sites of transcription are required to quantify engaged Pol II complexes at individual functional genomic regions. Libraries of nascent transcripts can be prepared with different techniques (reviewed in Wissink *et al.*, 2019). We use and recommend nascent transcription data obtained with Precision Run-On sequencing (PRO-seq; Kwak *et al.*, 2013), which marks active sites of transcription and isolates nascent transcripts from other cellular RNAs (Figure 2A). With certain limitations and caution (Wissink *et al.*, 2019; Wang *et al.*, 2019), the protocol reported here can also be applied to data derived from other strand-specific techniques, such as Global Run-On sequencing (GRO-seq; Core *et al.*, 2008) and mammalian NET-seq (Mayer *et al.*, 2015; Nojima *et al.*, 2015). Techniques that detect transcription initiation, such as Start-seq (Nechaev *et al.*, 2010) and PRO-cap (Mahat *et al.*, 2016a), can be used for mapping the functional genomic regions, but they do not allow quantification of engaged Pol II complexes at the distinct genomic regions.

**Figure 2.**
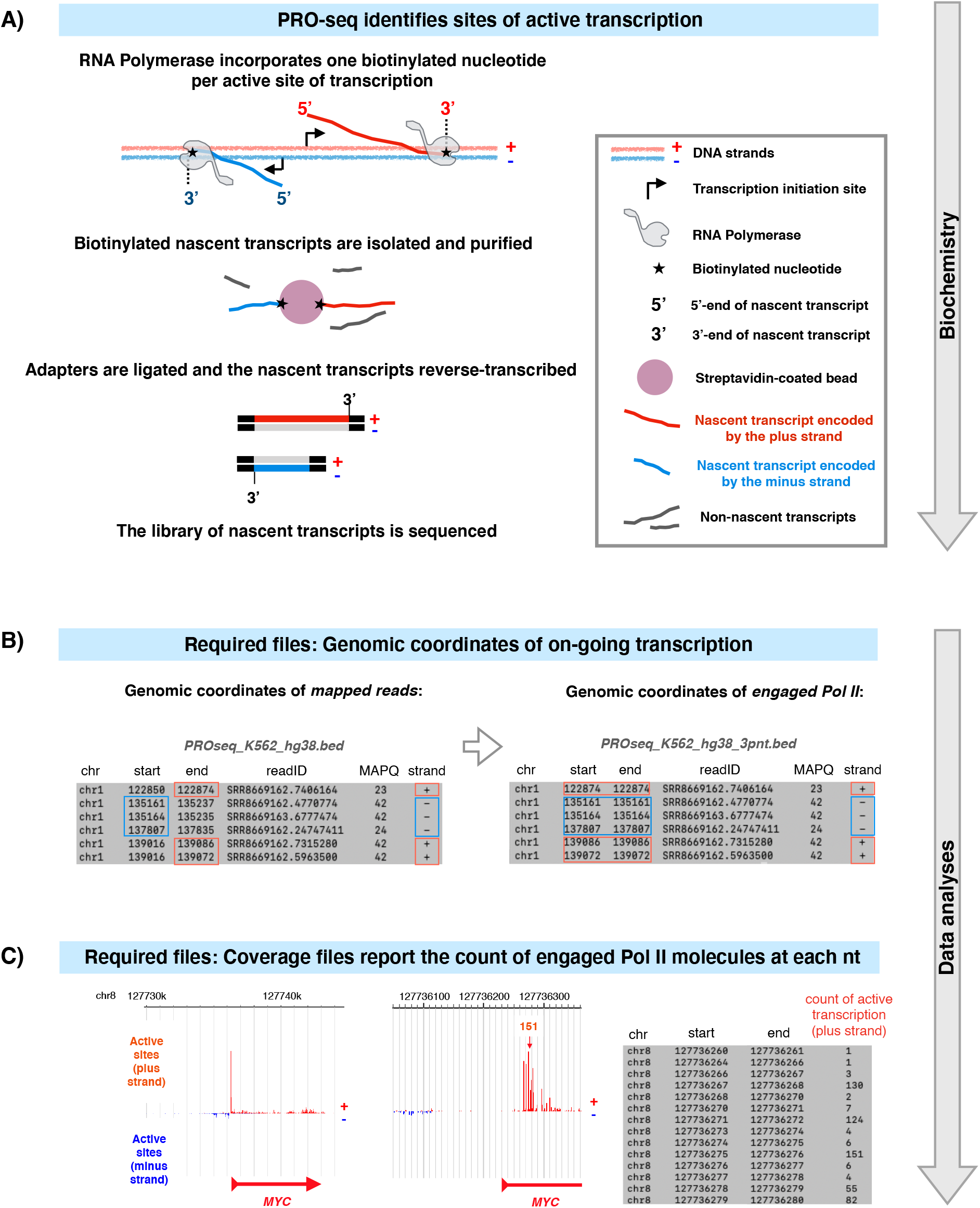
Generating nucleotide-resolution maps of nascent transcription. **A)** Schematic overview of PRO-seq library preparation. A biotinylated nucleotide is incorporated into each active site of transcription. The incorporated biotin-nucleotides allow isolation and purification of nascent transcripts. Adapters are ligated and the nascent transcripts reverse transcribed. Active sites of transcription are illustrated at two example reads encoded by the plus (red) and the minus (blue) strands. **B)** *Left panel:* Genomic coordinates of each mapped read (chromosome, start coordinate, end coordinate) are reported in a bed file. The coordinate representing the active site of transcription is indicated for two schematic reads, and boxed for each read in the example bed file. *Right panel:* The active sites of transcription are extracted from the coordinates of whole mapped reads and reported in a new bed file. **C)** Active sites of transcription are counted at each genomic coordinate. The resulting coverage files report the density of engaged transcription complexes at each nt. The *left panel* shows engaged transcription at *MYC*. The *middle panel* is an inset of the *MYC* promoter, and the *right panel* shows Pol II counts at individual nts of its promoter-proximal region.

### Required data format of nascent RNAs

Wet-lab protocols (Mahat *et al.*, 2016a) and computation pipelines (https://github.com/Vihervaara/PRO-seq-analyses; https://github.com/Danko-Lab/tutorials/blob/master/PRO-seq.md) for PRO-seq have been reported. Here, we begin with sequenced and mapped PRO-seq reads in bed format (Figure 2B) and strand-specific coverage files in bigWig format (Figure 2C). The bed format is a human-readable matrix that reports the genomic coordinates of each mapped read (single-end sequencing) or fragment (paired-end sequencing). Each row in the bed file lists one aligned entity (read or fragment) while the columns report its genomic coordinates (chromosome, start nucleotide, end nucleotide) and the strand that encodes the transcript (Figure 2B, left panel). For simplicity, we refer to the mapped entities as reads regardless of whether they were derived from single-end or paired-end sequencing. Coverage files include human-readable bedgraph (Figure 2C) and unreadable bigWig files where each genomic coordinate gets a value of intensity, reporting the count of transcription complexes at each nucleotide. Missing values equal to zero. We exemplify the protocol using PRO-seq data generated in human K562 cells (Vihervaara *et al.*, 2021), remapped against the latest hg38 reference genome. The bigWig and bed files, used here as input (see steps 1 and 13, respectively), can be directly obtained from Gene Expression Omnibus (GEO; https://www.ncbi.nlm.nih.gov/geo/) using accession code GSE181161.

### Downloading RefGene file with coordinates of gene transcripts

We derive functional genomic regions from the profile of active transcription and the refGene-annotated coordinates of gene transcripts. The refGene list of transcripts can be downloaded from the UCSC genome golden path: http://hgdownload.soe.ucsc.edu/goldenPath/. Here, human hg38 file (2020-08-17) is downloaded in step 4 using wget (https://www.gnu.org/software/wget/).

### De novo identification of active enhancers

Enhancers can be identified *de novo* using the pattern of divergent transcription (Danko et al, 2015; Azofeifa and Dowell, 2017; Vihervaara *et al.*, 2017; Chu *et al.*, 2018, Wang *et al.*, 2019). In this protocol, active promoters and enhancers are identified using discriminative Regulatory-Element detection from Global run-on sequencing (dREG; Wang *et al.*, 2019). dREG inputs unnormalized bigWig files of PRO-seq data and outputs coordinates of divergent transcription. dREG can be used either *via* web interface (https://django.dreg.scigap.org/) or installed locally (https://github.com/Danko-Lab/dREG). For broad accessibility, the web-based tool is used here. Unnormalized bigWig files are generally provided with PRO-seq data. If needed, the code below converts a bed file reporting the whole read (Figure 2B, left panel) into two 3’ coverage bedgraph files (Figure 2C). One of the bedgraph files reports active sites of transcription at the plus strand, the other at the minus strand. The bedgraph files are further processed to human unreadable bigWig format with bedgraphToBigWig tool (https://www.encodeproject.org/software/bedgraphtobigwig/). The required chrSizes.txt file is a two-column data frame that contains the chromosome names and the sizes (in nucleotides, nts), obtainable for the appropriate genome from http://hgdownload.cse.ucsc.edu/goldenpath/.

~~~
awk ‘$6 ⩵ “+”’ PROseq.bed | genomeCoverageBed −i stdin −3 −bg −g chrSizes.txt > PROseq_pl.bedgraph
awk ‘$6 ⩵ “-”’ PROseq.bed | genomeCoverageBed −i stdin −3 −bg −g chrSizes.txt > PROseq_temp.bedgraph
awk ‘{$4=$4*−1; print}’ PROseq_temp.bedgraph > PROseq_mn.bedgraph
bedgraphToBigWig PROseq_pl.bedgraph chrSizes.txt PROseq_pl.bigWig
bedgraphToBigWig PROseq_mn.bedgraph chrSizes.txt PROseq_mn.bigWig
~~~

### Analyzing genomic distribution of transcriptional regulators

Each step of transcription is coordinated by regulatory proteins. After identifying the functional genomic regions (Figure 3A-B) and counting engaged Pol II molecules at each region (Figure 3C-D), the protocol analyzes distribution of selected transcriptional regulators across the functional genomic regions (Figure 3E). Generating datasets for transcription factor binding sites is beyond the scope of this protocol. However, multiple techniques have been established to study protein and RNA localization to the genome, and raw data and peak coordinates for many chromatin-associating proteins can be obtained from databases such as ENCODE, modENCODE and GEO. We exemplify the localization of transcriptional regulators and chromatin marks using chromatin immunoprecipitation (ChIP)-seq data provide by the ENCODE (Consortium EP, 2011). The factors and histone marks analyzed here are TATA Box Binding Protein (TBP) and General Transcription Factor 2B (GTF2B), which are components of the Pre Initiation Complex, cell type-specific factors GATA1 and GATA2, Cohesin Complex Component RAD21, architectural protein CTCF, histone acetyl transferase p300 and histone mark H3K36me3 (Graphical Abstract and Figure 4). Steps to download the raw data and generate peak summits are found in github (https://github.com/Vihervaara/ChIP-seq_analyses).

**Figure 3.**
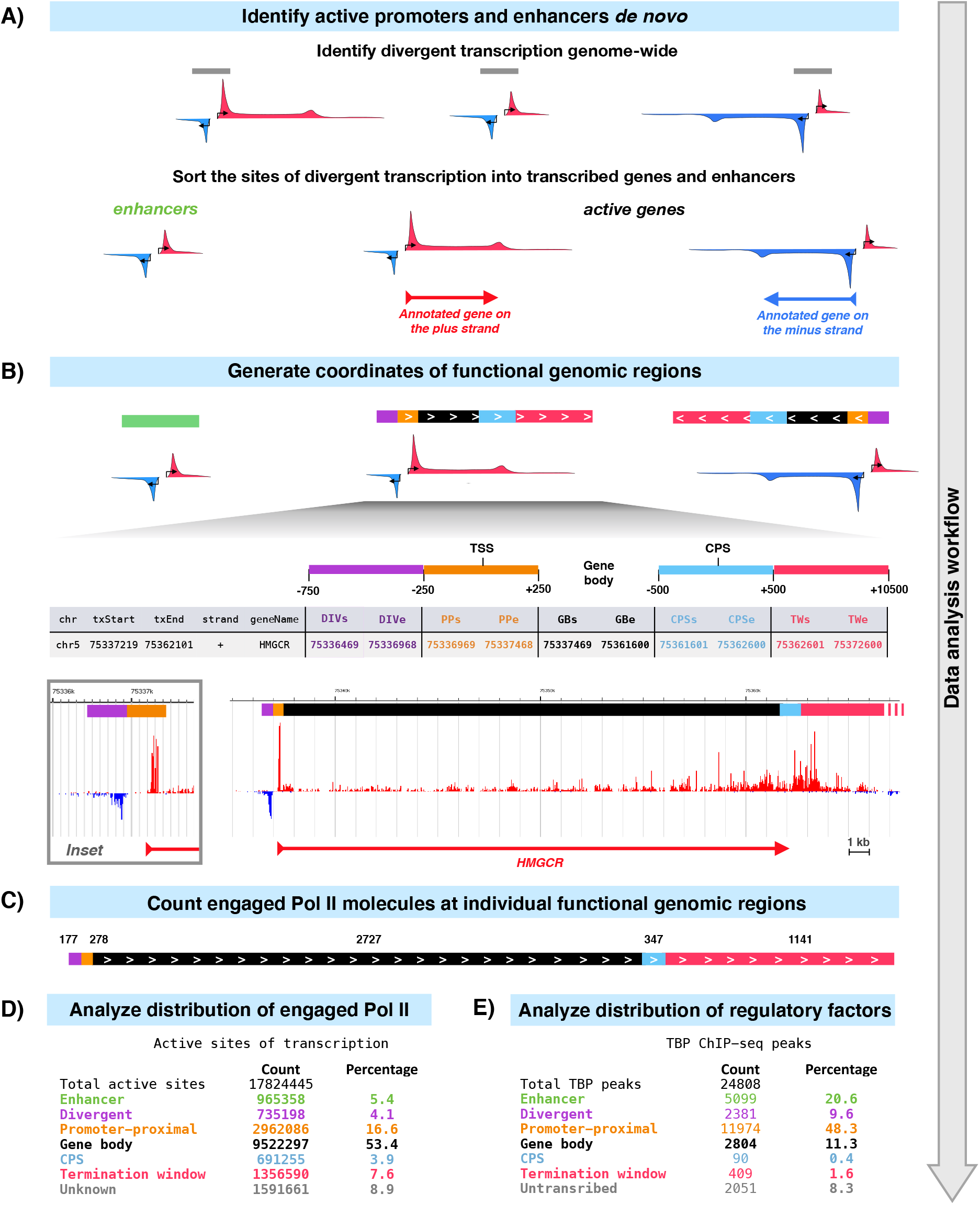
From identification of functional genomic regions to analyses of nascent transcription and transcription regulation. **A)** Schematic illustrations of the pattern of engaged Pol II at genes and enhancers. Sites of divergent transcription (grey bars) are identified *de novo* using dREG (Wang *et al.*, 2019). Divergent transcription that does not overlap with any annotated TSS of any gene is counted as an enhancer. Annotated genes with divergent transcription at the TSS are actively transcribed genes. **B)** At an enhancer, divergent transcription defines the enhancer coordinates (green bar). Active genes are divided into: upstream divergent transcription (−750 to −250 nt from the TSS; purple bar), promoter-proximal region (−249 to +250 nt from the TSS; orange bar), gene body (+251 from the TSS to −500 from the CPS; black bar), CPS (−499 to +500 nt from the CPS; light blue bar) and termination window (+501 to +10500 nt from the CPS). **C)** Engaged transcription complexes are counted at each identified functional genomic region. The count can be a raw count in the given library (exemplified here) or a normalized count that enable comparison between libraries (not shown). **D)** The count of engaged Pol II complexes and **E)** transcription factor binding sites uncovers the distribution of nascent transcription and its regulators across the genome. See the Graphical Abstract for illustration of the percentages in **D** and **E**. TBP: TATA box-binding protein. The TBP ChIP-seq data is from Consortium EP (2011).

**Figure 4.**
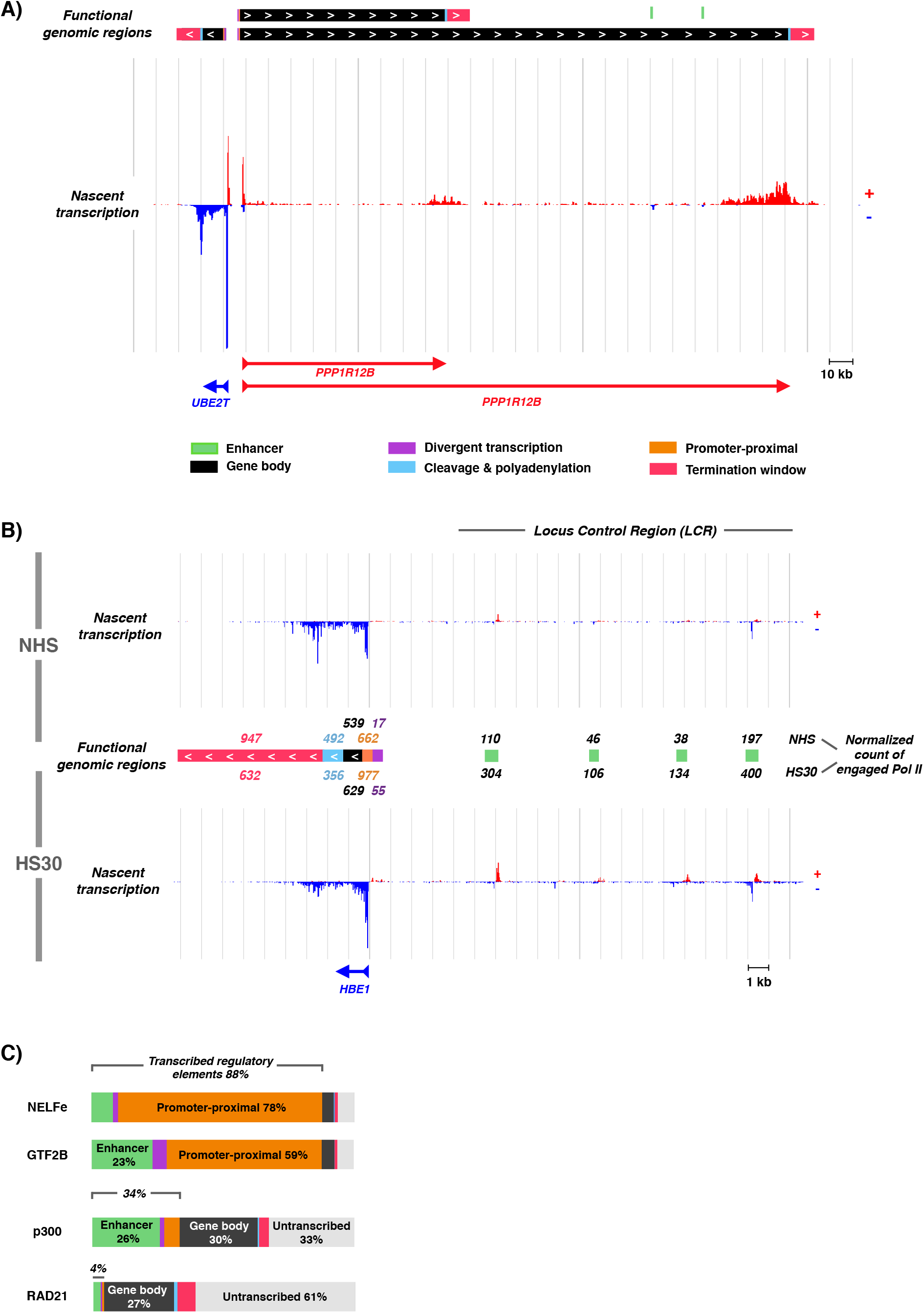
Expected results include A) identification of active genes and enhancers, B) quantification of nascent transcription complexes at functional genomic regions, and C) tracking genomic distribution of transcription regulators. **A)** Functional genomic regions identified at the *UBE2T* and *PPP1R12B* genes. **B)** Counts of nascent transcription complexes at hemoglobin beta 1 (*HBE1*) gene and the locus control region (LCR) of beta globin genes. The identified enhancers (green bars) represent the four functionally verified enhancers at the LCR in K562 cells. **C)** Localisation of selected chromatin-associating factors across the functional genomic regions. The brackets above the bars indicate the proportion of the factor peaks at transcribed regulatory elements (enhancers, upstream divergent transcription and promoter-proximal regions). The color coding of the bar is as in A), the additional gray regions indicates regions that were nor called as genes or enhancers (untranscribed / unknown). NELFe: negative elongation factor, subunit e. GTF2B: general transcription factor 2B. p300: histone acetyl transferase p300. RAD21: RAD21 Cohesin Complex Component. ChIP-seq data in figure C is from Consortium EP (2011).

### Obtaining data-analytical tools

The data-analyses reported here are conducted in command line environment of Apple OS X operating system. Lists of genomic coordinates are processed in R (https://www.r-project.org) and coordinates of functional genomic regions are compared to active sites of nascent transcription using bedtools (Quinlan et al, 2010; https://bedtools.readthedocs.io/en/latest/).

### Key Resources Table

| REAGENT or RESOURCE | SOURCE | IDENTIFIER |
| --- | --- | --- |
| Deposited data |  |  |
| PRO-seq | Raw data Vihervaara *et al.*, 2021. Remapping to hg38 in this study. | Raw data: GSE127844.<br>Remapped files: GSE181161 |
| TBP ChIP-seq raw data | Consortium EP, 2011 | https://www.ncbi.nlm.nih.gov/sra?term=SRX150574 |
| GTF2B ChIP-seq raw data | Consortium EP, 2011 | https://www.ncbi.nlm.nih.gov/sra/?term=SRR502177 |
| CTCF ChIP-seq raw data | Consortium EP, 2011 | https://www.ncbi.nlm.nih.gov/sra?term=SRX150487 |
| RAD21 ChIP-seq raw data | Consortium EP, 2011 | https://www.ncbi.nlm.nih.gov/sra/?term=SRR502023 |
| H3K36me3 ChIP-seq raw data | Consortium EP, 2011 | https://www.ncbi.nlm.nih.gov/sra?term=SRX067465 |
| p300 ChIP-seq raw data | Consortium EP, 2011 | https://www.ncbi.nlm.nih.gov/sra/?term=SRR502192 |
| NELFe ChIP-seq raw data | Consortium EP, 2011 | https://www.ncbi.nlm.nih.gov/sra/?term=SRR502172 |
| GATA1 ChIP-seq raw data | Consortium EP, 2011 | https://www.ncbi.nlm.nih.gov/sra?term=SRX186613 |
| Input ChIP-seq raw data | Consortium EP, 2011 | https://www.ncbi.nlm.nih.gov/sra?term=SRX150443 |
| Software and algorithms |  |  |
| Bedtools | Quinlan *et al.*, 2010 | v2.29.2 |
| Sra tools (fasterq-dump) | https://ncbi.github.io/sra-tools/fastq-dump.html | 2.8.0 |
| R | https://www.r-project.org | 4.1.0 |
| dREG | Wang *et al.*, 2019 | https://django.dreg.scigap.org/ |
| bedGraphToBigWig | https://www.encodeproject.org/software/bedgraphtobigwig/ | v 4 |
| IGV | Thorvaldsdóttir *et al.*, 2013 | http://software.broadinstitute.org/software/igv/ |
| kbro | Hojoong Kwak, Cornell University, NY, USA. | 2011 |
| Other |  |  |
| Human hg38 refGene file | UCSC downloads | http://hgdownload.soe.ucsc.edu/goldenPath/hg38/database/refGene.txt.gz |

## Step-by-Step Method Details

### Outline of the protocol

This protocol begins with identification of transcribed regulatory elements (regions initiating divergent transcription) *de novo* (Figure 3A). Subsequently, the coordinates of transcribed regulatory elements are intersected with annotated TSSs of gene transcripts (Figure 3A). Sites of divergent transcription that do overlap with TSSs of annotated genes indicate actively transcribed genes in the investigated condition. Sites of divergent transcription that do not occur in the vicinity of any annotated TSS of a gene are distal Transcribed Regulatory Elements (dTREs), also called enhancer candidates. We and others have previously verified that enhancer candidates identified from PRO-seq data contain marks of active enhancers, loop to gene promoters, and capture functionally verified enhancers (see Vihervaara *et al.*, 2017, 2021; Wang et al, 2019). For simplicity, we hereon refer to the enhancer candidates as enhancers.

At enhancers, the site of divergent transcription serves as an enhancer’s coordinates. Active genes, instead, are further divided into distinct regions (Figure 3B). A browser-compatible bed file is generated to visualize every functional genomic region, color-code each region according to the functional category, and display the count of engaged Pol II molecules at the region (Figure 3C). The count obtained here is the raw count of 3′-nts of PRO-seq reads (active sites) in the given dataset. This raw count can be normalized to allow comparison between datasets and account for different sequencing depths. Finally, distribution of engaged Pol II (Figure 3D) and selected regulatory proteins (Figure 3E) across the functional genomic regions is analyzed. For simplicity, the pipeline reported here uses a single condition of human K562 cells cultured in optimal growth condition.

### Run dREG to localize transcribed enhancers and active promoters

**Timing: [1-3h]**

1. Create an account at https://django.dreg.scigap.org/. Log in.

a. Choose dREG peak calling
b. Upload the unnormalized 3’-coverage bigWig *plus* strand file to the correct box
c. Upload the unnormalized 3’-coverage bigWig *minus* strand file to the correct box Please note that the bigWig files need to be unnormalized (minimum value +1 for the *plus strand* and −1 for the *minus strand*).
d. Choose a prefix (here ‘K562_hg38’) to describe your run and press Launch. The run time depends on the size of the file and available processing capacity, commonly ranging 1-3h.
2. When the run is complete, download the prefix.dREG.peak.full.bed.gz file and gunzip it.
3. Move the downloaded file to the working directory and rename it to a simpler form:

~~~
mv ~/Downloads/K562_hg38.dREG.peak.full.bed ~/pathToWorkingDirectory/dREGcalls_hg38_K562.bed
~~~

### Generate coordinates of functional regions

**Timing: [30 min]**

4. Download RefGene datafile, gunzip it and open in R.

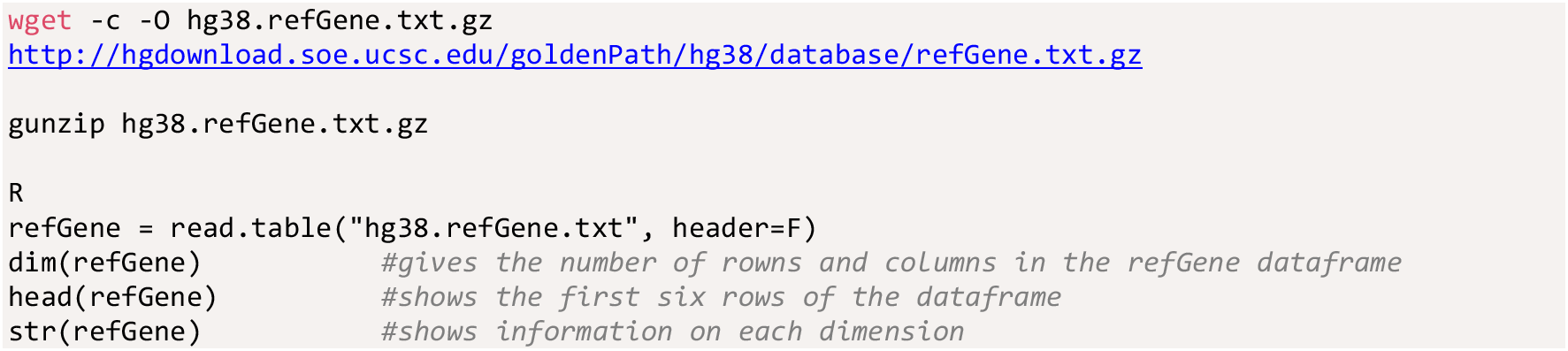
5. Name the refGene columns and remove unnecessary columns and chromosome entries.

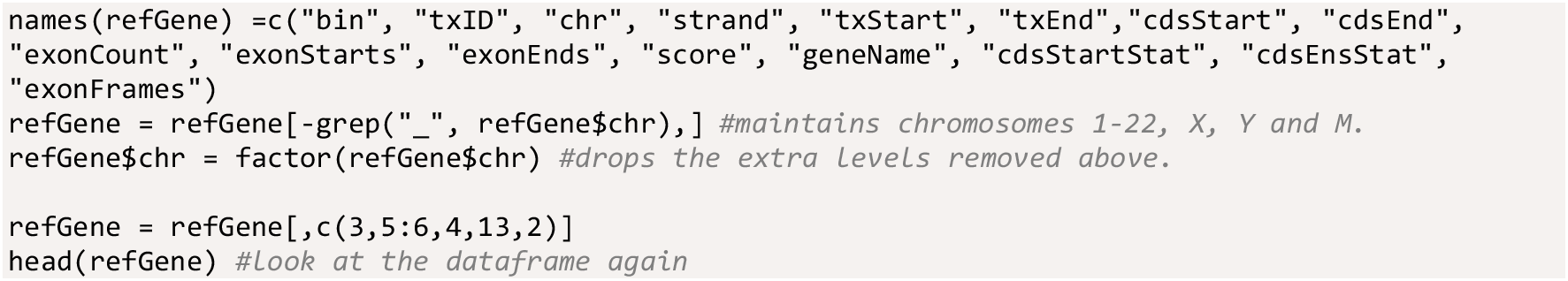
6. Generate coordinates of functional regions for every annotated gene transcript. In the RefGene data file, the ‘txStart’ is smaller than the same transcript’s ‘txEnd’. This is convenient when working with the coordinates of genes, but it leads to different columns reporting different functional sites depending on whether the gene is encoded by the plus or the minus strand. For example, the annotated TSS for genes on the plus strand is reported as txStart, while the TSS for genes on the minus strand is reported as txEnd. In the following steps, we add a column ‘TSS’ that reports the annotated TSS for each transcript. We then output a file that reports a 1000-nt window around the TSS. These windows of 1000 nt are intersected with the coordinates of dREG-identified sites of divergent transcription in step 7 to identify genes with active transcription and distal sites of divergent transcription, i.e. enhancers. We also obtain transcript-specific coordinates of promoter-proximal region (PP), divergent transcription (DIV), gene body (GB), CPS, and termination window (TW) according to the scheme in Figure 3B.

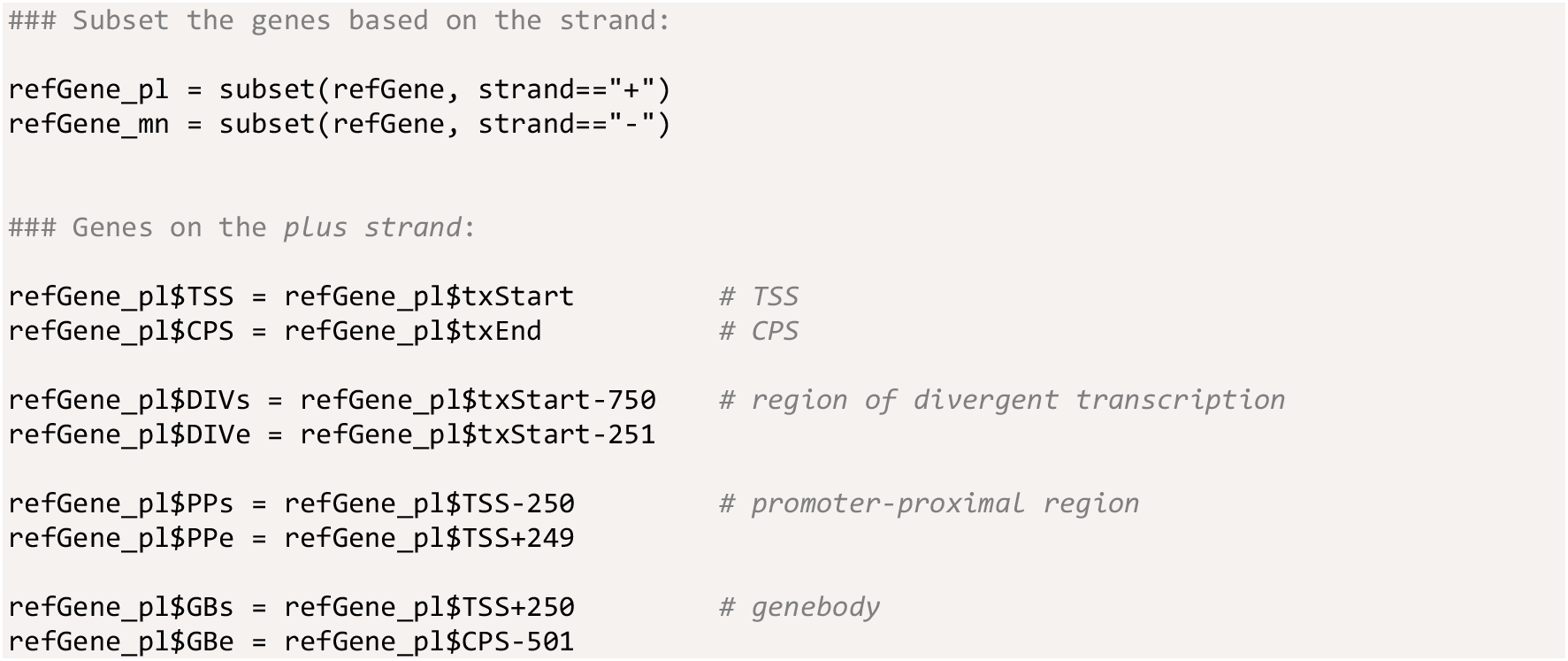

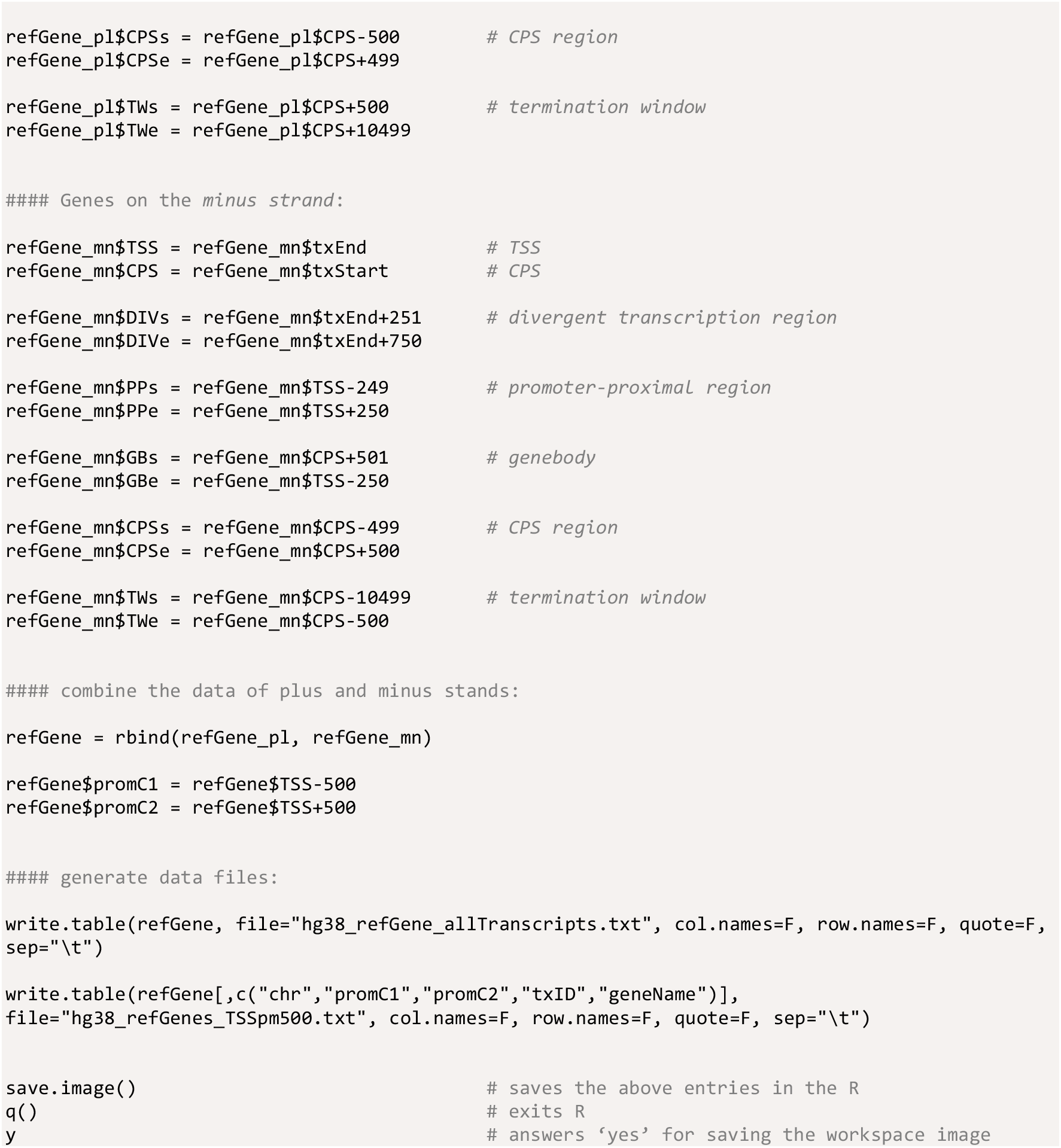

### Identify active promoters and transcribed enhancers

**Timing: [10 min]**

7. Intersect the dREG-identified regulatory regions with the 1000-nt window around TSSs.

a. Identify transcribed enhancers. These regulatory elements do not localize to any annotated TSS of a gene in the genome. The option −v directs the bedtools to output only entries in the file −a that do not show any overlap with coordinates given in file −b. The generated file contains the coordinates of transcribed enhancers.

~~~
bedtools intersect −v −a dREGcalls_hg38_K562.bed −b hg38_refGenes_TSSpm500.txt > enhancers.bed
~~~
b. Identify annotated genes with divergent transcription at the promoter. These are genes that have transcriptional activity in the investigated condition. The argument -wa maintains the entries in -a that have any overlap with -b. The argument -u controls that each entry in -a is written only once even if multiple dREG peaks would overlap with the same promoter. The generated file contains coordinates of gene TSSs that have transcriptional activity.

~~~
bedtools intersect -u -wa -a hg38_refGenes_TSSpm500.txt -b dREGcalls_hg38_K562.bed > activeGenes_hg38_K562.bed
~~~

### Write files of functional genomic coordinates

**Timing: [10 min]**

8. Return to R and read in the file with the TSSs of transcribed genes

~~~
R
Active = read.table(“activeGenes_hg38_K562.bed”)
~~~
9. Generate a new data frame in R that contains only actively transcribed genes. In essence, the ‘refGene’ data frame generated in step 6 is reduced here to contain only gene transcripts which initiate transcription. The ‘txID’ column contains an individual identification code for each transcript variant.

~~~
refGeneAct = subset(refGene, txID %in% Active[,4])
~~~
10. Write files that contain the coordinates of promoter-proximal and divergent transcription regions. These coordinates were generated in step 6.

~~~
write.table(refGeneAct[,c(“chr”,“PPs”,“PPe”,“geneName”,“txID”,“strand”)], file=“ppPolII.txt”,
col.names=F, row.names=F, quote=F, sep=“\t”)
write.table(refGeneAct[,c(“chr”,“DIVs”,“DIVe”,“geneName”,“txID”,“strand”)], file=“divTx.txt”,
col.names=F, row.names=F, quote=F, sep=“\t”)
~~~
11. Remove short genes before generating files with the gene body coordinates. This stage is needed to omit gene transcripts, where, due to shortness of the gene, the gene body would overlap with the promoter-proximal region (stretching to +500 from TSS) and CPS (starting from −500 from CPS) windows.

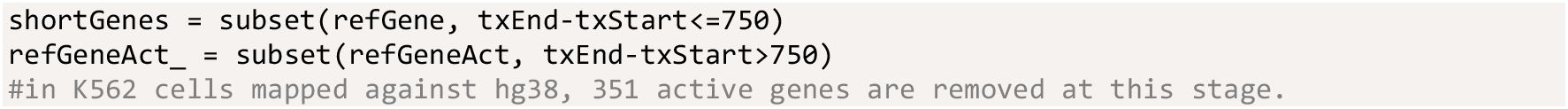
12. Write the files for the CPSs, gene body coordinates and transcription windows.

~~~
write.table(refGeneAct_[,c(“chr”,“CPSs”,“CPSe”,“geneName”,“txID”,“strand”)], file=“CPS.txt”,
col.names=F, row.names=F, quote=F, sep=“\t”)
write.table(refGeneAct_[,c(“chr”,“TWs”,“TWe”,“geneName”,“txID”,“strand”)], file=“TW.txt”,
col.names=F, row.names=F, quote=F, sep=“\t”)
~~~

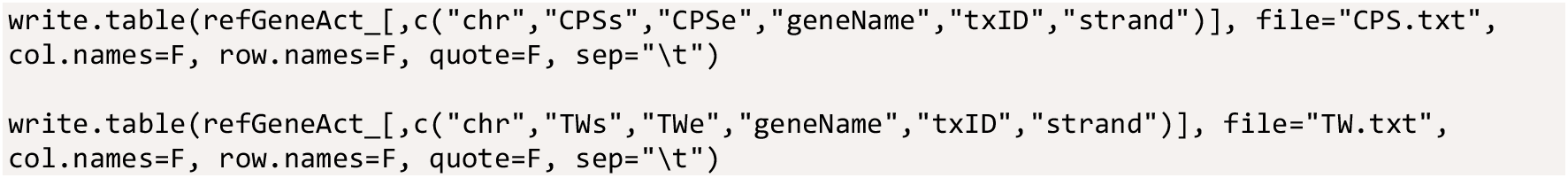

**Of note**: In RefGen for hg38 file, short genes constitute of 3,575 transcripts. Of these, only 351 genes were uniquely mappable, identified as ‘Active’ and, therefore, removed from our list of active gene transcripts in K562 cells.

### Allocate nascent transcription complexes to functional genomic regions

**Timing: [20 min]**

13. Generate a bed file that only contain the 3’-nucleotide (active site of transcription) of each read (Figure 2B). In the generated 3pnt.bed file (right panel in Figure 2B), each RNA Polymerase position is reported as an individual row.

a. Split the file based on the strand of the mapped read

~~~
echo retaining 3prime most coordinate of the bed file
awk ‘$6 ⩵ “+”’ PROseq_K562_hg38.bed > tempPL.bed
awk ‘$6 ⩵ “−”’ PROseq_K562_hg38.bed > tempMN.bed
~~~
b. Active site of transcription (3’-most nt) is the coordinate given in the third column of plus strand reads and the second column for minus strand reads. In this step, the active site of transcription (single nt) will be placed both to the second and the third column.

~~~
awk ‘{$2 = $3; print}’ tempPL.bed > tempPL_3p.bed
awk ‘{$3 = $2; print}’ tempMN.bed > tempMN_3p.bed
~~~
c. The reads from plus and minus strands are combined, the data is converted to tab-delimited, and the reads sorted based on genomic coordinates. Intermediary files are removed.

~~~
cat tempPL_3p.bed tempMN_3p.bed | tr ‘ ‘ ‘\t’ > temp_3p.bed
sortBed -i temp_3p.bed > PROseq_K562_hg38_3pnt.bed
rm *temp*
~~~
14. Intersect the coordinates of nascent transcription with the coordinates of functional genomic regions.

The coordinates of genomic regions were saved above as individual.txt files. Here, the number of engaged Pol II molecules (rows in the 3pnt.bed file) are allocated to distinct functional categories.

**Note:** To allocate each engaged Pol II complex only once, we use -u option in the bedtools intersect command and sequentially allocate the coordinates of active transcription to the distinct genomic categories. In each round, two different files are generated: File 1 retains active sites that *localize* to the given functional category (options -u and -wa). File 2 reports the active sites that *do not localize* to the given functional category (controlled with option -v). The file 2 will be used in the subsequent round to ensure that each engaged Pol II is allocated only to one genomic region. The order of the intersections is: i) promoter-proximal regions, ii) sites of divergent transcription, iii) enhancers, iv) CPSs, v) gene bodies, and vi) termination windows. In this order, promoter-associated Pol II molecules are not counted into enhancer transcription. Furthermore, transcription at intragenic enhancers is allocated to enhancers instead of gene bodies. Finally, termination windows can be relatively short or extend over several kilobases (Vilborg *et al.*, 2017). In this strategy, long (10 kb) termination windows are queried, but only Pol II complexes that do not overlap with any other functional genomic region are allocated to a gene’s termination window.

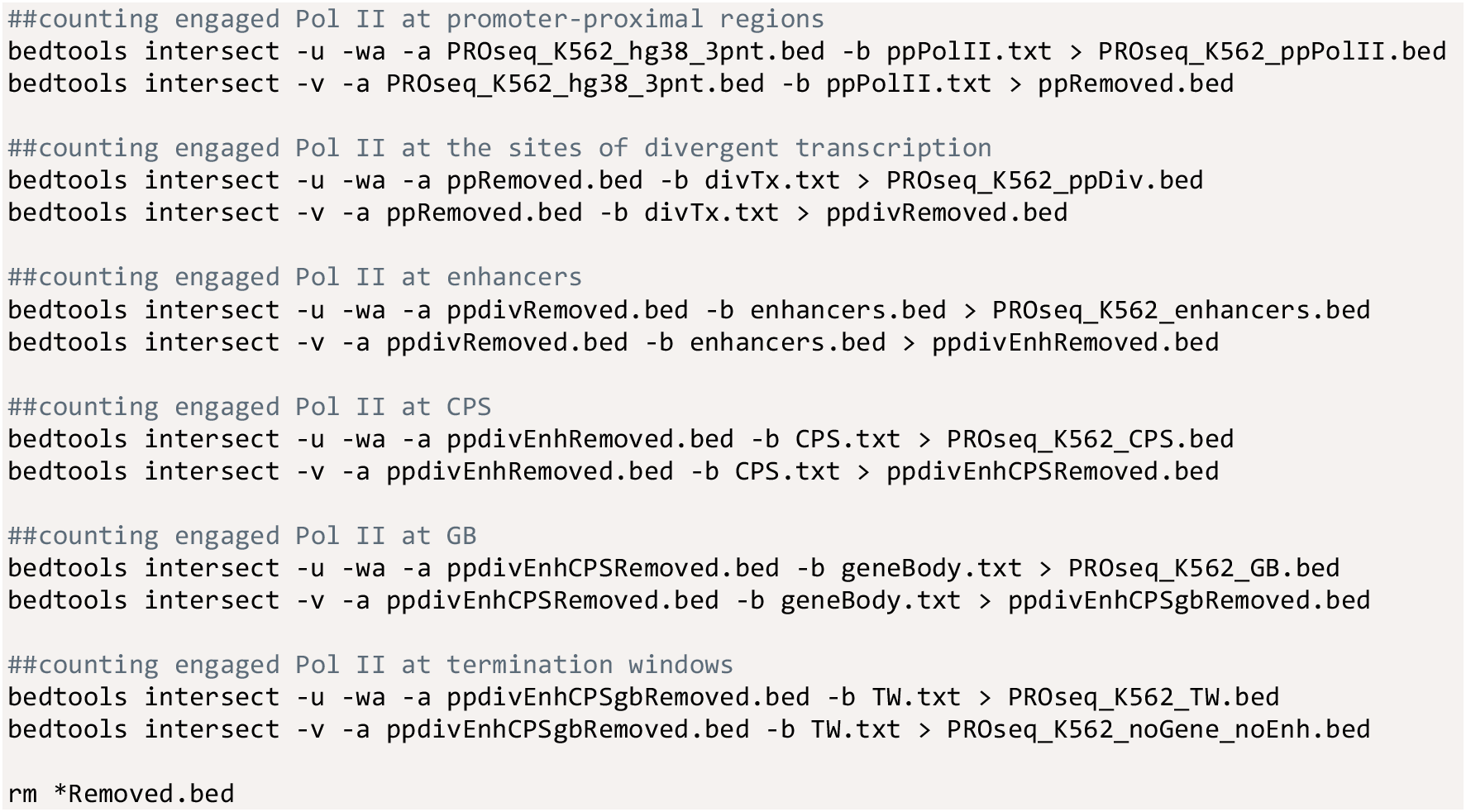

### Quantify distribution of nascent transcription across functional genomic regions

**Timing: [10 min]**

15. Plot the counts of engaged Pol II molecules at distinct categories of functional genomic regions.

a. Initiate a script that collects counts of engaged Pol II molecules at the distinct genomic regions. The counts will be printed in the.txt file as well as on the terminal window.

~~~
script counts_at_functional_regions.txt
~~~
b. Plot the counts of rows in each intersected bed file. The number of rows in each file corresponds to the number of active sites of transcription in the given category of functional genomic regions.

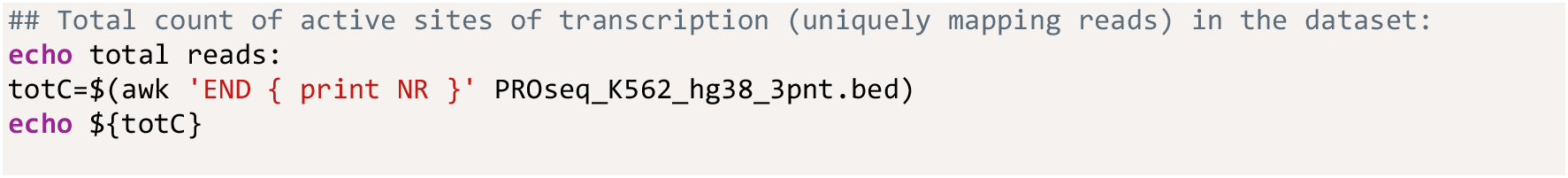

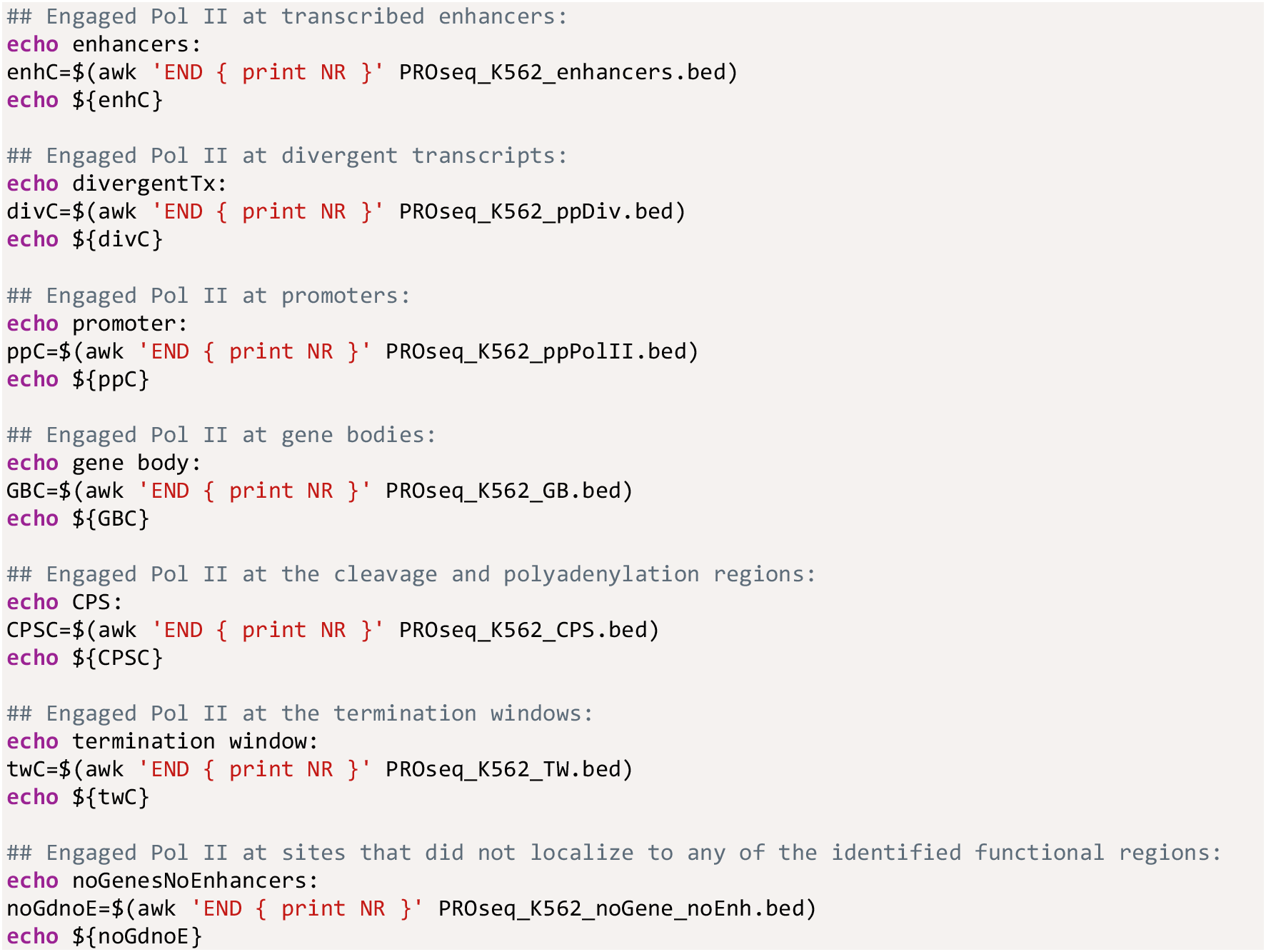
c. Terminate the log script by pressing <u>***control + D***</u> in the terminal window.

The file ‘counts_at_functional_regions.txt’ now reports the number of engaged Pol II molecules at each category of functional genomic regions. Figure 3D shows the counts in a table format and includes percentages of engaged Pol II in each category. The pie chart in the Graphical Abstract illustrates the distribution of engaged Pol II complexes across the functional genomic regions.

**NOTE:** The counts generated in this protocol correspond to sites of active transcription (3’-nt of a read) in the dataset. Here, we only have a single sample and, therefore, focus on the distribution (proportions) of active sites across the genomic regions. With PRO-seq data, multiple conditions can be compared by computing a normalization factor that accounts for differences in data handling and sequencing depth. Detailed description of normalization factors is out of the scope of this study. In brief, invariant whole-genome spike-in from a distinct organism can be added to all samples before the run-on reaction. This equal amount of foreign chromatin provides a count of nascent transcription against which the samples can be normalized (Booth *et al.*, 2018; Vihervaara et al,. 2021). Alternatively, genomic regions where no changes are detected during a time course can be utilized to generate a normalization factor between samples (Mahat *et al.*, 2016b; Vihervaara *et al.*, 2017).

### Quantify nascent transcription complexes at individual genomic regions

**Timing: [15 min]**

16. Intersect coordinates of the active sites of transcription with the coordinates of individual functional genomic regions.

~~~
bedtools intersect -c -wa -a ppPolII.txt -b PROseq_K562_ppPolII.bed > ppPolCounts.tmp
bedtools intersect -c -wa -a divTx.txt -b PROseq_K562_ppDiv.bed > ppDivCounts.tmp
bedtools intersect -c -wa -a enhancers.bed -b PROseq_K562_enhancers.bed > enhancerCounts.tmp
bedtools intersect -c -wa -a geneBody.txt -b PROseq_K562_GB.bed > geneBodyCounts.tmp
bedtools intersect -c -wa -a CPS.txt -b PROseq_K562_CPS.bed > CPSCounts.tmp
bedtools intersect -c -wa -a TW.txt -b PROseq_K562_TW.bed > TerminationWinCounts.tmp
~~~
17. Color code the distinct categories of functional genomic regions. The color coding is in bed-compatible rgb format (0-255,0-255,0-255).

~~~
awk -F ‘\t’ -v OFS=’\t’ ’{ $(NF+1) =“243,132,0”; print }’ ppPolCounts.tmp > ppPolCounts.bed
awk -F ‘\t’ -v OFS=’\t’ ‘{ $(NF+1) =“178,59,212”; print }’ ppDivCounts.tmp > ppDivCounts.bed
awk -F ‘\t’ -v OFS=’\t’ ‘{ $(NF+1) =“115,212,122”; print }’ enhancerCounts.tmp >
enhancerCounts.bed
awk -F ‘\t’ -v OFS=’\t’ ’{ $(NF+1) =“0,0,0”; print }’ geneBodyCounts.tmp > geneBodyCounts.bed
awk -F ‘\t’ -v OFS=’\t’ ‘{ $(NF+1) =“103,200,249”; print }’ CPSCounts.tmp > CPSCounts.bed
awk -F ‘\t’ -v OFS=’\t’ ‘{ $(NF+1) =“255,54,98”; print }’ TerminationWinCounts.tmp >
TerminationWinCounts.bed
~~~
18. Combine the files. Add an extra column “.” to obtain genome browser compatible bed file for visualization.

~~~
cat ppPolCounts.bed ppDivCounts.bed enhancerCounts.bed geneBodyCounts.bed CPSCounts.bed
TerminationWinCounts.bed > catRegions.temp
awk -F ‘\t’ -v OFS=’\t’ ‘{ $(NF+1) =“.”; print }’ catRegions.temp > catRegions2.temp
~~~
19. Reorganize the columns and add a header track.

~~~
awk ‘{print $1 “\t” $2 “\t” $3 “\t” $7 “\t” $5 “\t” $6 “\t” $2 “\t” $3 “\t” $8}’
catRegions2.temp > catRegions3.temp
awk ‘!seen[$1,$2,$3,$6]++’ catRegions3.temp | sortBed > catRegions4.temp
touch headerLine.txt
echo track name=“functional_genomic_regions” itemRgb=“On” ≫ headerLine.txt
cat headerLine.txt catRegions4.temp > functionalGenomicRegions.bed
rm *.temp
rm headerLine.txt
~~~

The generated file can be read into a genome browser to visualize the identified functional genomic regions, including the count of engaged Pol II at each region (Figures 3C and 4A-B).

NOTE: As mentioned above, the number of engaged Pol II complexes at each region is derived from the raw reads of the data. This count depends on the sequencing depth. To get a normalized count of engaged Pol II, the column 4 (reporting raw count) in the ‘functionalGenomicRegions.bed’ file can be multiplied with a normalization factor.

### Analyze the distribution of chromatin-associated factors to functional genomic regions

**Timing: [30 min if summit files are available, up to hours if they need to be generated from raw data]**

20. Count genome-associating factors at distinct functional genomic regions.

a. Obtain or generate datasets of interest. Place them in the folder of your working directory. Here, we use TBP, GATA1, CTCF, H3K36me3, NELFe, GTF2B, p300 and RAD21 ChIP-seq data generated by the ENCODE (Consortium EP, 2011). We remapped the ENCODE data to hg38 (https://github.com/Vihervaara/ChIP-seq_analyses). As starting material, please use a file that reports the summit coordinate of each peak, named with ‘_summits.bed’ ending, for example: K562_TBP_summits.bed, K562_GATA1_summits.bed, K562_CTCF_summits.bed K562_H3K36me3_summits.bed
b. The peak summits for each given chromatin-associated factor are intersected with the distinct genomic regions. The strategy described in step 14 is used to ensure that each identified enrichment of a factor at the genome (peak) is counted once. For efficiency, we use a loop function that takes one bed file of mapped peak summits at a time. To define which datasets are analyzed in the loop function, please place names of factors in quotation marks, separated by a space, after the *for x in* code below. In the loop, the *${x}* will be replaced with the factor name, one listed factor after another.
c. Run the code below using the factors of your choice. Please, ensure that the files are placed in the correct folder (working directory) and that the file names correspond to the names in the code. The files listing the coordinates of functional genomic regions were generated in steps 1-12.

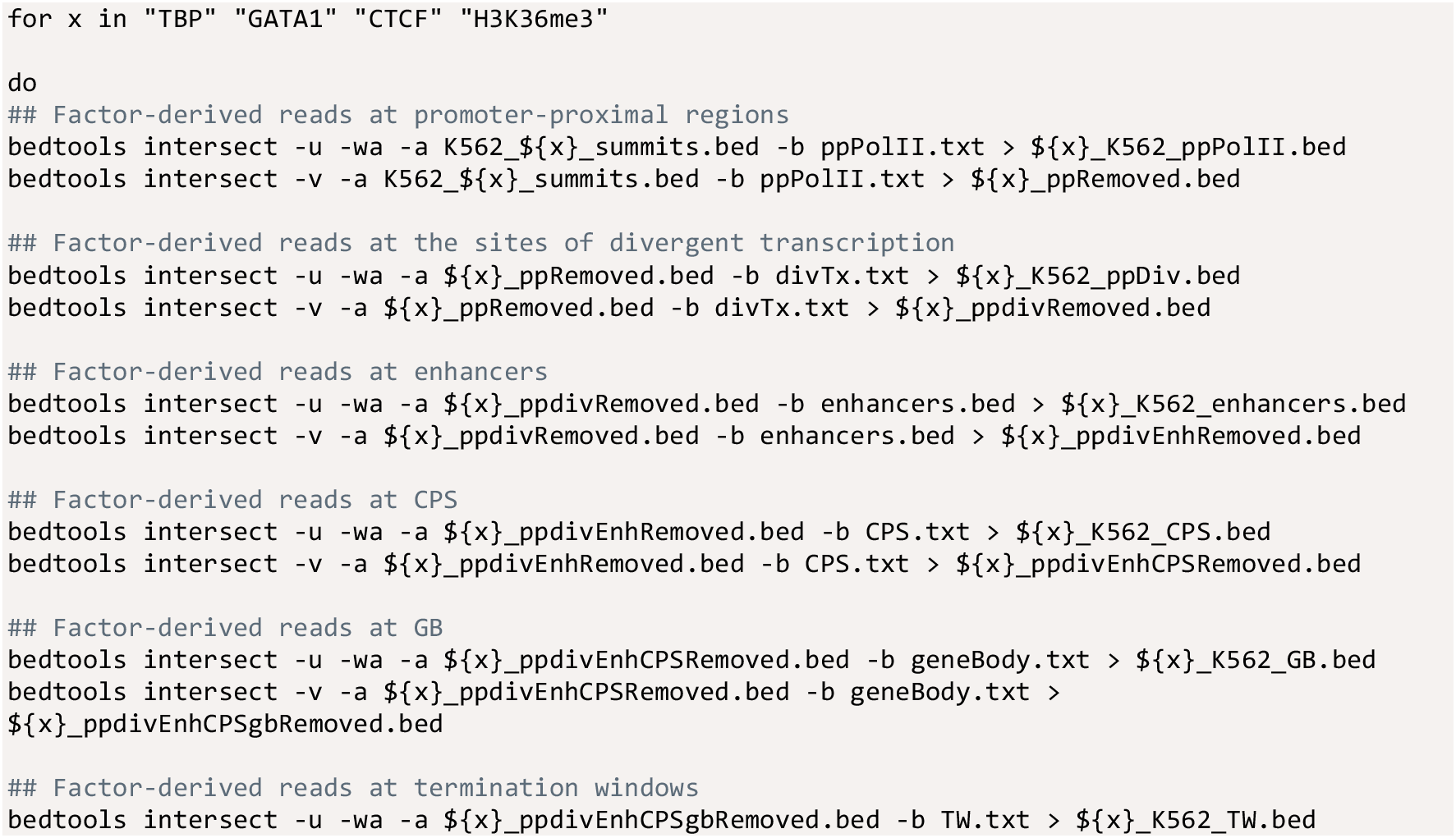

~~~
bedtools intersect -v -a ${x}_ppdivEnhCPSgbRemoved.bed -b TW.txt > ${x}_K562_noGene_noEnh.bed
rm *Removed.bed done
~~~
21. Plot the counts of chromatin-associated factors at distinct categories of functional regions.

a. Initiate the script that collects factor counts at the functional genomic regions.

~~~
script Factor_counts_at_functional_regions.txt
~~~
b. Plot the counts of rows in each intersected bed file.

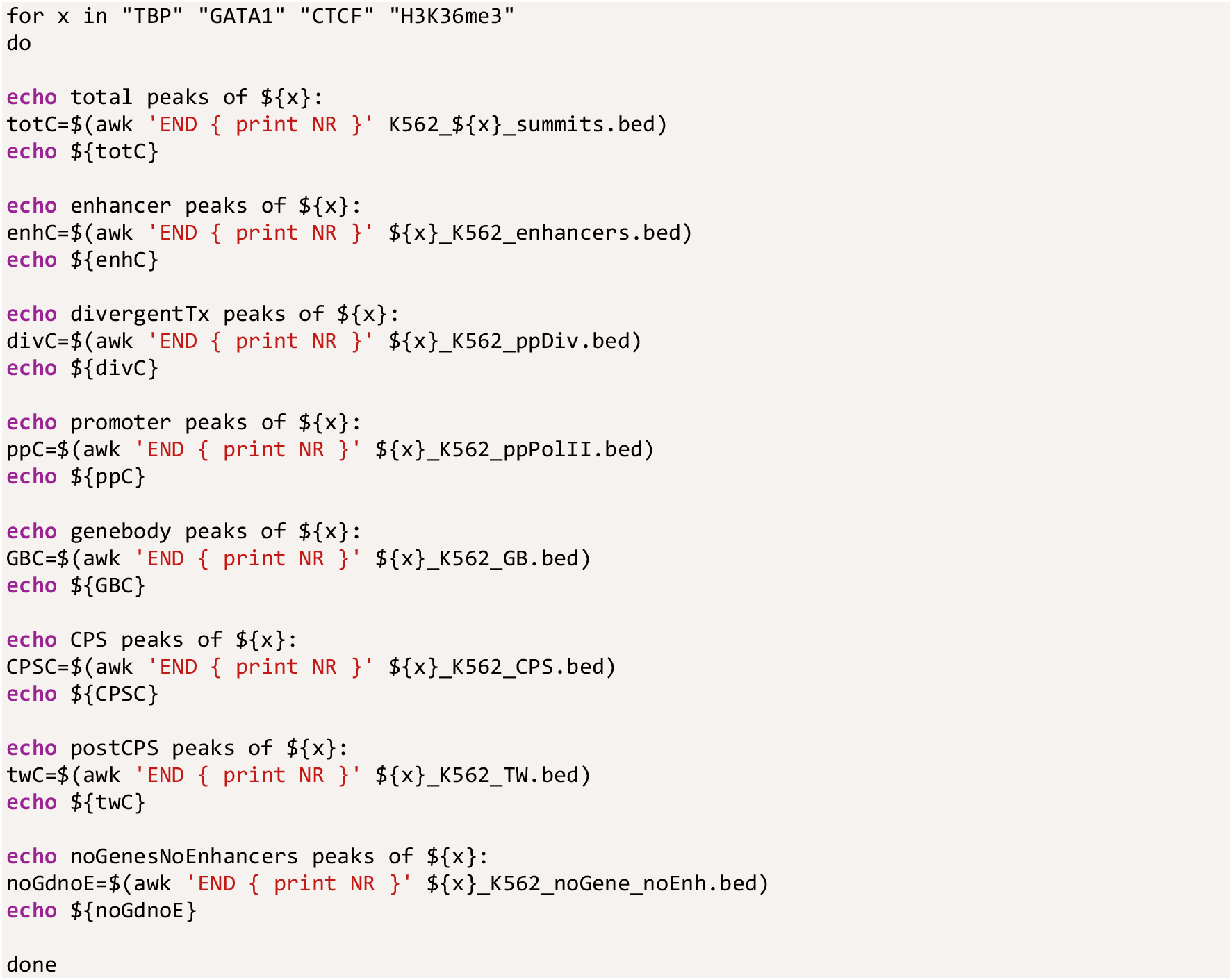
c. Terminate the log script by pressing <u>***control + D***</u> in the terminal window.
22. The file ‘Factor_peaks_at_functional_regions.txt’ reports the number of ChIP-seq peaks at each functional category. Figure 3E exemplifies the counts for TBP peaks in a table format, including the percentages of binding sites at each category of functional genomic regions. The bar charts in the Graphical Abstract illustrates the distribution of TBP, GATA1, CTCF and H3K36me3 across the functional genomic regions. In Figure 4E, bar charts for additional factors (NELFe, GTF2B, p300 and RAD21) are included.

## Expected Outcomes

### Counting nascent RNA synthesis at functional genomic regions

This protocol inputs active sites of transcription, and outputs coordinates of transcribed genomic regions. A file that lists and visualizes individual functional genomic regions is generated in steps 1-19. In this *functional_genomic_regions.bed* file, each block represents the coordinates of a functional genomic region, and the block color indicates the category of the region (green: enhancer; purple: divergent transcription; orange: promoter-proximal region; black: gene body; light blue: cleavage and polyA site; pink: termination window). Figure 4A shows transcribed isoforms and functional genomic regions of *UBET2* and *PPP1R12B* genes, as well as two intergenic enhancer candidates on the longer isoform of *PPP1R12B*. The protocol also counts engaged Pol II complexes at each identified functional genomic region. The raw count of active sites of transcription is given in column 4 of the *functional_genomic_regions.bed* file, appearing above the color-coded block when visualized in a genome browser (Figure 4B). The raw count can be multiplied with a normalization factor to allow comparison of distinct PRO-seq datasets and/or account for sequencing depth. In figure 4B, the count of engaged transcription machineries is exemplified at the hemoglobin beta locus, spanning from *HBE1* gene to the region’s upstream locus control element (LCR). In PRO-seq used here (Vihervaara *et al.*, 2021), the same amount of *Drosophila* S2 cells were spiked into each human K562 sample as a means for normalization. Figure 4B shows the raw count of reads in untreated (NHS) K562 cells above the identified functional genomic regions. The PRO-seq sample of 30-minute heat shocked cells (HS30) has been normalized against the NHS by comparing the spike-in reads (normalization factor 0.94 derived from: 36,872 spike-in reads in NHS / 39,407 spike-in reads in HS30). The counts can be further normalized to take into account the sequencing depth. Comparing engaged Pol II complexes at functional genomic regions allows analyzing changes in the distribution of nascent transcription at individual genomic regions. As an example, the hemoglobin beta locus (Figure 4B) demonstrates heat-induced accumulation of transcription machineries at promoter-proximal regions and enhancers.

### Identifying chromatin-associated factors at transcribed genomic regions

Transcriptional steps occur at specific functional genomic regions, and they are coordinated by transcription factors, co-factors, elongation factors, chromatin remodelers, histone marks, architectural proteins, RNA-processing factors, and regulatory RNA. Subsequently, tracking interactions at the genome is essential for understanding the regulatory logic of transcription. In steps 20-22 of the protocol, ChIP-seq data from the ENCODE (Consortium EP, 2011) is utilized to map binding sites of chromatin-associating proteins across the functional genomic regions. The summit coordinate of each ChIP-seq peak, representing the highest point of a factor enrichment, is queried against the coordinates of functional genomic regions. The code identifies genomic regions bound by each queried factor and outputs the distribution of each factor’s binding sites across the functional genomic regions (Graphical Abstract and Figure 4C). TBP and GTF2B, both components of the Pre-Initiation Complex, localize primarily to promoter-proximal regions but they are also abundant at enhancers and sites of divergent transcription (Graphical Abstract and Figure 4C). GATA1 and p300, instead, bind more often enhancers than promoters, and architectural protein CTCF and RAD21 have most of their enrichments in untranscribed regions (Graphical Abstract and Figure 4C). Histone mark H3K36me3 displays a striking 98% of its binding sites at gene bodies (93%), downstream CPSs (2%) or termination windows (3%, Graphical Abstract). These patterns of factor localizations reflect the steps of transcription they coordinate and can reveal regulatory stages at individual genomic regions.

## Limitations

This protocol is limited to genomes where active genes and enhancers initiate transcription to diverging orientations. Consequently, to identify active genes and enhancers *de novo* this protocol requires strand-specific mapping of nascent transcription. Furthermore, precise positional information of active sites of transcription is essential for correct quantification of engaged Pol II complexes at the identified functional genomic regions. A good quality and adequate sequencing depth of nascent transcription data is also required. With shallow sequencing, pause-peaks are poorly visible and divergent transcription becomes less evident, particularly at weakly transcribed genes and enhancers. In human genome, >15 million uniquely mapping reads derived from >5 million cells, provides a robust identification of divergently transcribed genes and enhancers. This protocol can be run with a lower sequencing depth, but please note that weakly transcribed genes and enhancers might not be called as active regulatory elements. Likewise, low coverage, e.g. due to small amount of cells as starting material, can impair the sensitivity for calling functional genomic regions at weakly transcribed parts of the genome.

## Troubleshooting

### Problem 1

Terminal shell gives a ‘file not found’ argument.

### Potential Solution

Check your working directory by typing ‘pwd’ in the terminal shell. If the files you are attempting to use are not in your working directory, change the working directory by typing ‘cd ‘ and dragging the correct folder after it in the terminal shell window. Ensure also that all the files you are attempting to use are in the working directory and that the names of the files exactly match the names given in the code. For example, the ‘K562_’ in the file names provided here need to be removed or replaced when analyzing PRO-seq data from another cell type.

### Problem 2

Terminal shell does not recognize the tools or packages.

### Potential Solution

Install correct software. In Mac OS, bedtools and wget can be installed for example *via* homebrew (https://brew.sh):

~~~
/bin/bash -c “$(curl -fsSL https://raw.githubusercontent.com/Homebrew/install/HEAD/install.sh)”
brew install bedtools
brew install wget
~~~

For installing R, see https://www.r-project.org for instructions to download the correct R.app. R, wget and bedtools can be run *via* terminal shell.

### Problem 3

dREG finds only a small number of sites with divergent transcription.

### Potential Solution

Ensure that unnormalized bigWig files are uploaded in dREG. Ensure also that the PRO-seq library is of adequate quality and has a good sequencing depth. If the nascent RNA-seq library has a low count of uniquely mapped reads or low complexity, the Pol II pause peaks might not become visible. In these cases, the pattern of divergent transcription might not be adequate for calling active genes and transcribed enhancers.

### Problem 4

Only a small number of active genes is identified.

### Potential Solution

The genomes of human and mouse are are well-annotated. In other species, a large fraction of the genes might remain to be identified. If working with a poorly annotated genome, the active genes are likely counted as enhancers since the reference genome does not report a gene TSS at the site of divergent transcription. As a possible solution, check that the latest version of gene annotation is used. Furthermore, tools such as dREG.HD (Chu *et al.*, 2018) and primaryTranscriptAnnotation in R (Anderson *et al.*, 2020) could predict whether the transcription initiates from a gene or an enhancer, and identify the length of the transcript unit, respectively.

## Resource Availability

### Lead Contact

Further information and requests for resources and reagents should be directed to and will be fulfilled by the Lead Contact, Anniina Vihervaara.

### Materials Availability

This study did not generate new unique reagents.

### Data and Code Availability

The PRO-seq data utilized in this study is described in Vihervaara *et al.*, (2021) and accessible *via* GEO (GSE127844). The input data files used here were remapped to hg38 genome and are available *via* GEO accession code GSE181161. The ChIP-seq datasets were generated by the ENCODE (Consortium EP, 2011), the raw data of which is available *via* accession code GSE31477. The code to generate input files used here have been deposited to github (https://github.com/Vihervaara/PRO-seq-analyses; https://github.com/Vihervaara/ChIP-seq_analyses).

## Acknowledgments

This work was financially supported by Science for Life Laboratory (A.V.), Svenska Tekniska Vetenskapsakademi i Finland (A.V.), The Sigrid Jusélius Foundation (A.V. and L.S.), Academy of Finland (A.V. and L.S.), Åbo Akademi University (L.S.).

## Author Contributions

A.R., S.C. and A.V. conducted the data-analyses. S.C. and A.V. prepared the figures, L.S. and A.V. conceived and designed the work. All the authors wrote and/or edited the manuscript.

